# Tomato *CRABS CLAW* paralogues interact with chromatin remodelling factors to mediate carpel development and floral determinacy

**DOI:** 10.1101/2021.08.19.456989

**Authors:** Laura Castañeda, Estela Giménez, Benito Pineda, Begoña García-Sogo, Ana Ortiz-Atienza, Rosa Micol-Ponce, Trinidad Angosto, Juan Capel, Vicente Moreno, Fernando J. Yuste-Lisbona, Rafael Lozano

## Abstract

*CRABS CLAW* (*CRC*) orthologues play a crucial role in floral meristem (FM) determinacy and gynoecium formation across angiosperms, key developmental processes for ensuring successful plant reproduction and crop production. Here, we revealed that the incomplete penetrance and variable expressivity of the carpel-inside-carpel phenotype observed in flowers of the tomato *fruit iterative growth* (*fig*) mutant is due to a lack of function of a homologue of the *CRC* gene, *Solanum lycopersicum CRCa* (*SlCRCa*). Likewise, a comprehensive functional analysis of *SlCRCa* and *SlCRCb* paralogues, including Arabidopsis complementation experiments, allowed us to propose that they operate as positive regulators of FM determinacy by acting in a compensatory and partially redundant manner to safeguard the proper formation of flowers and fruits. Furthermore, we provide the first evidence for the role of putative *CRC* orthologues as members of the chromatin remodelling complex that terminates floral stem cell activity by repressing *WUSCHEL* expression.

## Introduction

Angiosperms are characterised by producing flowers as reproductive organs to ensure their effective reproduction. From their outermost to the innermost whorls, flowers typically consist of sepals, petals, stamens and carpels that are sequentially generated from a pool of stem cells located in the floral meristems (FM) (Krizek and Fletcher, 2005; Aichinger et al., 2012). Once a set number of floral organs has been initiated, stem cell activity is arrested, and the FM is thereby determined to form the female reproductive structure known as gynoecium. The precise timing of this developmental event, also referred to as floral determinacy, is a pivotal process that establishes a defined number of floral organs arising from the FM (Sablowski, 2007; Sun et al., 2009; Sun and Ito, 2015).

In the model species *Arabidopsis thaliana*, the homeodomain transcription factor WUSCHEL (WUS) is responsible for the maintenance of the stem cell activity in the FM. The MADS-box transcription factor AGAMOUS (AG) plays a key role in the timing regulation of FM termination by repressing *WUS* expression (Laux et al., 1996; Mayer et al., 1998; Liu et al., 2011). AG turns off the stem cell maintenance programme involving transcriptional repression of *WUS* by different pathways; directly, by a mechanism that implicates chromatin remodelling and the recruitment of the Polycomb Group (PcG) protein TERMINAL FLOWER2/LIKE HETEROCHROMATIN PROTEIN1 (TFL2/LHP1) at the *WUS* locus (Liu et al., 2011; Guo et al., 2018), and indirectly through the transcriptional induction of two key downstream targets, *KNUCKLES* (*KNU*) and *CRABS CLAW* (*CRC*), which act through independent pathways to synergistically regulate *WUS* repression and ensure an adequate FM determination process (Payne et al., 2004; Sun et al., 2009, 2014, 2019; Yamaguchi et al., 2017, 2018). The *KNU* gene encodes a C2H2 zinc-finger protein whose expression is activated by AG, a process that requires a time-delay induction regulated by epigenetic modification of histones at the *KNU* locus (Sun et al., 2014). Once induced, KNU binds to the *WUS* promoter, which causes the eviction of SPLAYED (SYD), a chromatin remodelling factor required for *WUS* activation, and mediates the subsequent deposition of H3K27me3 for stable Polycomb-mediated repression of *WUS* (Kwon et al., 2005; Sun et al., 2019). Furthermore, AG also positively regulates *MINI ZINC FINGER2* (*MIF2*) expression during flower development, which acts as an adaptor protein to form a transcriptional repressor complex together with KNU, the transcriptional corepressor TOPLESS (TPL), and the chromatin remodelling protein HISTONE DEACETYLASE19 (HDA19). Within this complex, MIF2 binds to the *WUS* locus, leading to an epigenetically repression of *WUS* expression through histone deacetylation (Kagale and Rozwadowski, 2011; Krogan et al., 2012; Bollier et al., 2018). Concurrently, the YABBY transcription factor CRC, a direct target of AG, mediates auxin homeostasis and establishes auxin maxima during carpel primordia initiation by repressing *TORNADO2* (*TRN2*) and upregulating the auxin synthesis gene *YUCCA4* (*YUC4*). The proper auxin maxima mediated by *CRC* contributes to the termination of FM cells proliferation through *WUS* repression and triggers the subsequent gynoecium formation (Yamaguchi et al., 2017, 2018).

In tomato (*Solanum lycopersicum* L.), the molecular mechanisms underlying *SlWUS* transcriptional regulation during floral development are of agronomic interest, as mutations leading to a spatial and temporal expansion of its expression domains in FMs result in flowers with extra carpels, which give rise to larger multilocular fruits (Muños et al., 2011; van der Knaap et al., 2014; Xu et al., 2015; Rodríguez-Leal et al., 2017; Yuste-lisbona et al., 2020). Likewise, alterations in *SlWUS* transcriptional repression during flower development potentially lead to indeterminate fruits, which makes floral determinacy a developmental process closely related to fruit shape, an important fruit quality attribute as it influences consumer’s acceptance and postharvest handling. Recently, Bollier et al. (2018) have proposed a conserved molecular mechanism regulating FM determinacy in Arabidopsis and tomato. Thus, the interaction of tomato MIF2 and KNU orthologues, INHIBITOR OF MERISTEM ACTIVITY (SlIMA) and SlKNU, allows for the recruitment of tomato TPL and HDA19 orthologues, SlTPL1 and SlHDA1, to form a transcriptional repressor complex at the *SlWUS* locus. In the present study, we provide novel insights into the genetic and molecular mechanism involved in FM determinacy and carpel development. Through combining mapping-by-sequencing, RNA interference (RNAi) and CRISPR/Cas9 techniques, we revealed that the carpel-inside-carpel phenotype observed in *fruit iterative growth* (*fig*) mutant plants is due to the lack of function of the *S. lycopersicum CRC* homologue *SlCRCa*. Furthermore, we carried out a detailed functional analysis of tomato *CRC* paralogues, *SlCRCa* and *SlCRCb*, which allowed us to uncover for the first time the role of putative *CRC* orthologues as members of the molecular regulatory network that epigenetically represses *WUS* through histone deacetylation to ensure the proper termination of floral stem cell activity.

## Results

### The *fig* mutation impairs carpel determinacy

The *fruit iterative growth* (*fig*) mutant was isolated from the screening of a collection of T_1_ segregating T-DNA lines generated from the tomato cultivar P73 (Pérez-Martín et al., 2017). A detailed phenotypic analysis revealed that the vegetative development of *fig* mutant plants was indistinguishable from that in wild-type (WT) ones, whereas significant differences were observed during flower and fruit development (Fig. 1a-c and Supplementary Table 1). *fig* flowers developed an elevated number of organs in all whorls, being this increase more evident in carpels. Thus, *fig* ovaries are composed of numerous carpels that lead to anomalous fruits which show secondary fruit structures growing in an indeterminate way that appeared from the inside of the principal fruit (Fig. 1a-c and Supplementary Table 1).

**Fig. 1.**
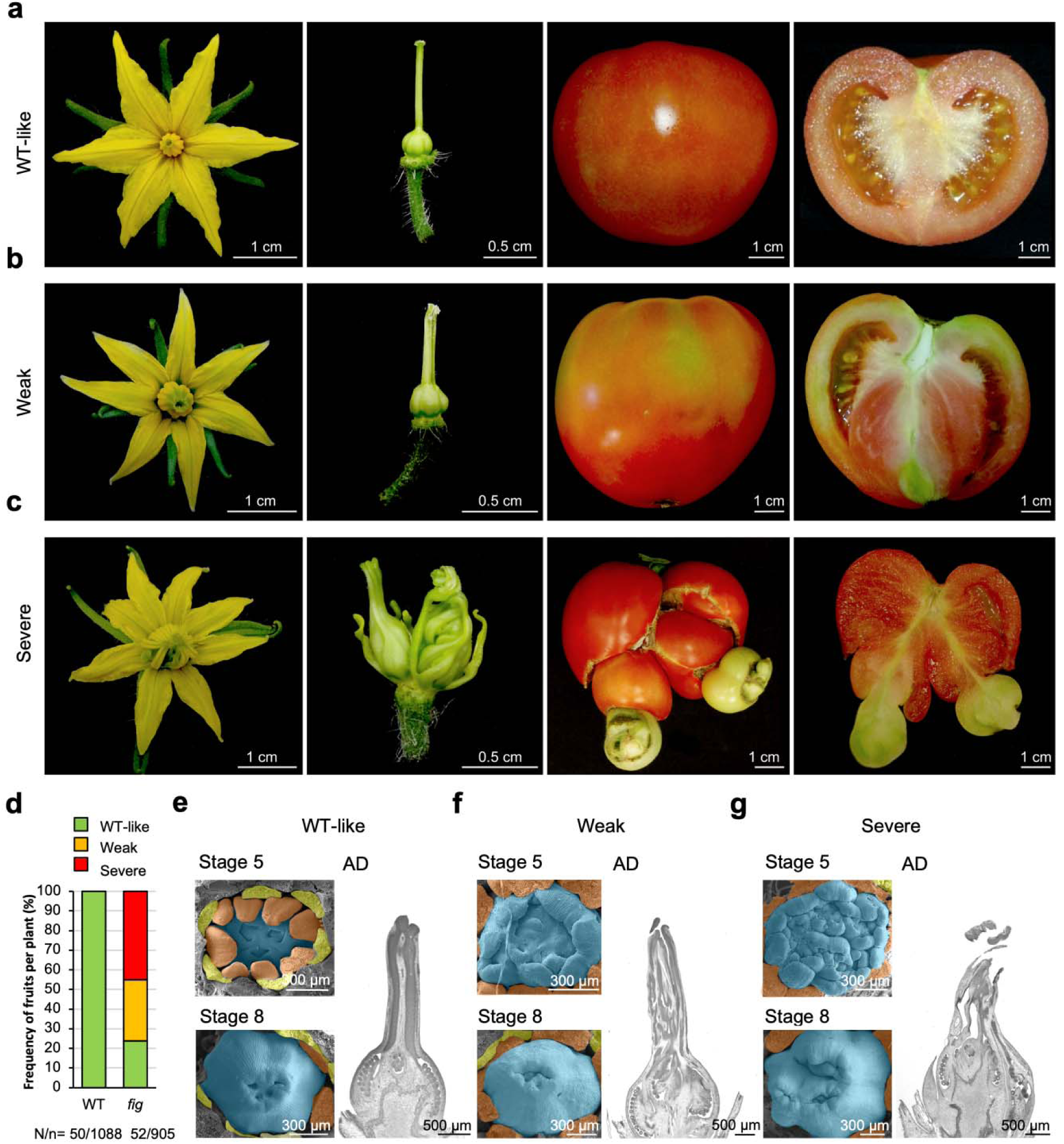
Phenotypic characterisation of the *fig* mutant. **a-c**, Representative *fig* flowers, pistils, closed fruits, and longitudinal open fruits (from left to right) showing WT-like (identical to wild-type, WT) (**a**), weak (**b**) and severe (**c**) phenotypes. **d**, Percentage of different types of fruits (WT-like, weak or severe) produced by WT (cv. P73) and *fig* plants. N, number of plants evaluated; n, number of fruits harvested. **e-g**, Scanning electron microscopy images of flowers at stages 5 and 8 of floral development, and histological sections of flowers at anthesis day (AD) stage exhibiting WT-like (**e**), weak (**f**) and severe (**g**) phenotypes. Sepals were removed in samples at stage 5 whereas only the carpels were maintained in developing flowers at stage 8. Petals are coloured in yellow, stamens in orange, and carpels in blue.

The observed phenotypic segregation (39 WT: 15 *fig*) was consistent with a monogenic recessive inheritance of the *fig* mutation (X^2^ = 0.22; *P* = 0.64). However, the phenotype of *fig* plants showed incomplete penetrance and variable expressivity, as a gradation of phenotypes was displayed within the same mutant plant, even within the same inflorescence. Thereby, variable flower and fruit phenotypes were observed in *fig* plants, which were classified as WT-like (indistinguishable from WT), weak and severe indeterminate phenotypes (Fig. 1a-c and Supplementary Fig. 1), being the average production of fruits with said severe *fig* phenotype close to 50% per mutant plant (Fig. 1d).

A scanning electron microscopy (SEM) study in developing flowers revealed that the first visible anomalies could be detected at floral stage 5 as estimated by floral bud size (Brukhin et al., 2003). At this stage, the carpels emerge, and the ovary cavities become visible showing abnormal carpel structures in both weak and severe *fig* flowers (Fig. 1f,g). Later, at stage 8, WT pistils are formed by 3-4 carpels (Fig. 1e), whereas an increased number of carpels are observed in *fig* flowers resulting in incomplete fused pistils (Fig. 1f,g and Supplementary Fig. 1). Moreover, histological sections of flowers at anthesis day showed that *fig* pistils have shorter and thicker styles and are composed of numerous carpels that grow one inside another, which strongly suggests that *fig* mutation affects carpel determinacy. These differences were more accentuated in severe *fig* flowers (Fig. 1c,g), producing ovaries threefold bigger than the WT ones at anthesis day stage (Fig. 1a,e). As a result of this variable range of *fig* flower phenotypes, we could observe both weak *fig* fruits producing secondary fruit structures only visible inside the fruit (Fig. 1b), and severe *fig* fruits where these secondary fruit structures emerged from inside and were visible outside the fruit (Fig. 1c and Supplementary Fig. 1). Despite such abnormalities and although more extreme *fig* fruits produced fewer seeds, *fig* mutants gave rise to viable seeds.

### *FIG* encodes a homologue of the Arabidopsis *CRC* gene

Although the *fig* mutant was isolated from a T-DNA insertional mutant collection, molecular analysis performed by Southern blot and PCR assays revealed that the *fig* mutation was not associated with a T-DNA insertion (Supplementary Fig. 2), suggesting that somaclonal variation produced during the *in vitro* culture process is responsible for the mutant phenotype. To identify the causative mutation underlying the *fig* phenotype, we performed a mapping-by-sequencing strategy using an F_2_ mapping population derived from crossing *fig* with the wild tomato *S. pimpinellifolium* (accession LA1589). A total of 783 F_2_ plants were scored for ovary and fruit development, from which 212 plants produced fused carpels and indeterminate fruits. The phenotypic segregation observed in this interspecific F_2_ progeny (571 WT: 212 *fig*) was consistent with a monogenic recessive inheritance pattern of the *fig* mutation (χ^2^ = 1.80, *P* = 0.18). To perform mapping-by-sequencing, we conducted genome sequencing of two DNA pools containing 50 WT and 25 *fig* F_2_ plants with the most strongly indeterminate phenotype. Genome-wide analysis of the allele frequencies revealed a region encompassing the centromere of chromosome 1 (2.4-70 Mb) with a strong bias towards tomato reference alleles (Fig. 2a). The variant analysis encompassing the candidate genomic region led to the identification of three SNPs mapping at the fourth intron of the *Solyc01g010240* gene, which also showed a marked decrease in the sequence coverage of the mutant pool compared to other coding and non-coding regions of this gene. To confirm the polymorphisms identified, PCR and Sanger sequencing analyses with primers flanking the region containing these SNPs were performed in WT and *fig* plants. The results showed that the PCR product from *fig* genomic DNA, which included the fourth intron of the *Solyc01g010240* gene, not only contained the three SNPs mentioned above, but it was also larger than expected. Sequence analysis verified that a DNA fragment of 367 bp was inserted at position 5,053,687 on the chromosome 1 of the mutant genome (Assembly SL4.0), interrupting the intron sequence between the *Solyc01g010240* exons 4 and 5 (Fig. 2b). This insertion shares a sequence identity of 86% with the Long Terminal Repeat (LTR) copy placed between 6,336,872 and 6,337,201 positions on the chromosome 10 (Assembly SL4.0), which may have acted as a transposable element. Co-segregation analysis performed in the F_2_ segregating population showed that the 212 *fig* mutant plants were homozygous for the LTR insertion, whereas 393 and 178 phenotypically WT plants were hemizygous or lacked the 367 bp insertion, respectively. Hence, results indicate that the *fig* phenotype co-segregated with the LTR insertion at the fourth intron of the *Solyc01g010240* gene.

**Fig. 2.**
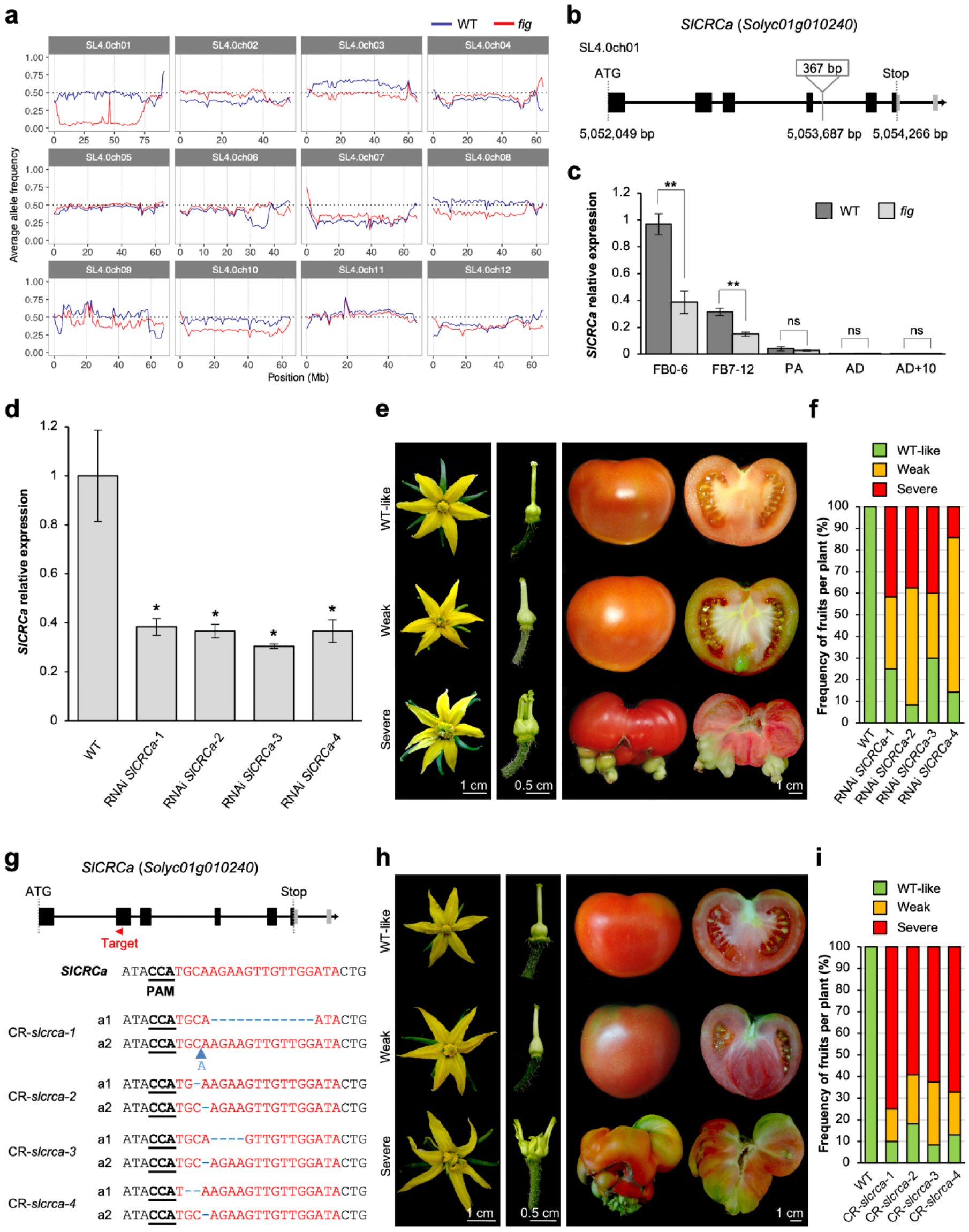
*FIG* encodes a homologue of the Arabidopsis *CRC* gene. **a**, Distribution of the average allele frequency of wild-type (WT; blue line) and *fig* (red line) pools grouped by chromosomes. **b**, Schematic representation of the *SlCRCa* gene (introns are represented by black lines, coding and UTRs are in black and grey boxes, respectively) and the 367 bp sequence inserted at the fourth intron in *fig* mutants. **c**, Relative expression of *SlCRCa* in WT and *fig* flowers at different stages of floral development. FB0-6, floral buds from stages 0 to 6; FB7-12, floral buds from stages 7 to 12; PA, flowers at pre-anthesis stage; AD, flowers at anthesis stage; AD+10, flowers 10 days after anthesis stage. **d**, *SlCRCa* transcripts quantification of flowers at stage FB0-6 from RNAi *SlCRCa* and WT lines. **e**, Representative RNAi *SlCRCa* flowers, pistils and fruits displaying WT-like, weak and severe mutant phenotypes. **f**, Percentage of different types of fruits harvested from WT and T_0_ RNAi *SlCRCa* plants. **g**, CRISPR/Cas9-*slcrca* (CR-*slcrca*) alleles identified by cloning and sequencing PCR products from the *SlCRCa* targeted region from four T_0_ CRISPR/Cas9 plants. Black bold and underlined letters indicate protospacer-adjacent motif (PAM) sequences, blue dashed lines show InDel mutations, blue letter and arrow indicate an insertion sequence. **h**, Representative CR-*slcrca* flowers, pistils and fruits exhibiting WT-like, weak and severe phenotypes. **i**, Percentage of different types of fruits harvested from WT and T_0_ CR-*slcrca* plants. In **c**,**d** data are means ± standard deviations (SD) of three biological and two technical replicates. A two-tailed, two-sample Student’s *t* test was performed, and significant differences are represented by asterisks: *, *P* < 0.01; **, *P* < 0.001; ns, no statistically significant differences.

Despite the fact that the 367 bp insertion occurs at a non-coding region, we considered the *Solyc01g010240* gene as a strong candidate to be responsible for the *fig* phenotype, since it encodes a homologue of the Arabidopsis *CRC* gene, *SlCRCa*, a putative transcription factor of the YABBY gene family that has been described as a key gene involved in the regulation of carpel and nectary development, as well as in FM determination (Alvarez and Smyth, 1999; Bowman and Smyth, 1999). To determine the effect of the LTR insertion on *SlCRCa* expression, cDNA cloning and qRT-PCR expression analyses were performed. Sequence analysis of WT and mutant cDNAs revealed that both were identical; however, mutant plants showed significantly reduced *SlCRCa* expression levels in floral buds at developmental stages 0-6 and 7-12 (Fig. 2c). Thus, the 367 bp insertion in the fourth intron of *SlCRCa* reduced its transcript level to >2-fold although still allowed the production of WT *SlCRCa* mRNA, suggesting that this intron may contain a transcriptional regulatory element critical for the maintenance of the *SlCRCa* spatio-temporal expression pattern.

To corroborate whether the lack of SlCRCa function is responsible for the *fig* phenotype, we generated RNAi-mediated knockdown lines with reduced levels of *SlCRCa* transcripts in the cultivar P73 genetic background. Four independent first-generation (T_0_) RNAi *SlCRCa* diploid lines were evaluated, which showed >2-fold decrease in *SlCRCa* expression in floral buds at stage 0-6 (Fig. 2d). These reduced *SlCRCa* transcript levels led to the development of flowers with supernumerary carpels that produced indeterminate fruits (Fig. 2e). As happened with the *fig* mutation, RNAi *SlCRCa* lines displayed a variable range of flower phenotypes, resulting in the development of fruits with WT-like, weak and severe indeterminate phenotypes (Fig. 2e,f). Furthermore, we engineered knockout mutations by using the CRISPR/Cas9 system with a single guide RNA targeting the second exon of the *SlCRCa* gene (Fig. 2g). We assessed four independent T_0_ diploid lines (CR-*slcrca*) that were biallelic for edited knockout alleles (Fig. 2g). The CR-*slcrca* lines developed flowers and fruits which undoubtedly resembled the phenotype observed in *fig* mutants, and they also produced a wide diversity of flower and indeterminate fruit phenotypes, most of them classified as severe (Fig. 2f,h). Thereby, although there were differences in the percentages of fruits with severe mutant phenotype, both knockdown (RNAi) and knockout (CRISPR/Cas9) alleles resulted in similar phenotypes with incomplete penetrance and variable expressivity, indicating that any deficiency in SlCRCa function might lead to a loss of FM determinacy. Taken together, all these results strongly support that the 367 bp insertion in the fourth intron of the *SlCRCa* gene is responsible for the alterations in carpel and fruit development observed in the *fig* mutant.

### *SlCRCa* expression is restricted to carpels

To better understand the function of *SlCRCa*, we monitored its expression pattern in several vegetative and reproductive tissues. As expected from the phenotype of *fig* mutant plants, *SlCRCa* expression was mainly restricted to flowers during early developmental stages. Thus, the higher expression level of *SlCRCa* was found in floral buds at stages 0-6 (Fig. 3a). We next performed an RNA *in situ* hybridisation analysis in young flowers to further examine the temporospatial expression patterns of *SlCRCa*. This analysis revealed that *SlCRCa* transcripts were accumulated specifically and uniformly when carpel primordia were initiated at floral stage 3 (Fig. 3b), and persisted in these primordia at floral stage 6, when carpels were growing up and the primordium of placenta emerged (Fig. 3c). At floral stage 8, *SlCRCa* expression was located at the adaxial surface on the base of the ovary walls, as well as in the most distal cells of the developing gynoecium which would give rise to the style and the stigma (Fig. 3d). However, *SlCRCa* mRNA was not detected in the ovary walls at floral stage 9, and its expression remained in the distal part of the gynoecium (Fig. 3e). At later stages of floral development, there was no evidence of detectable *SlCRCa* transcripts.

**Fig. 3.**
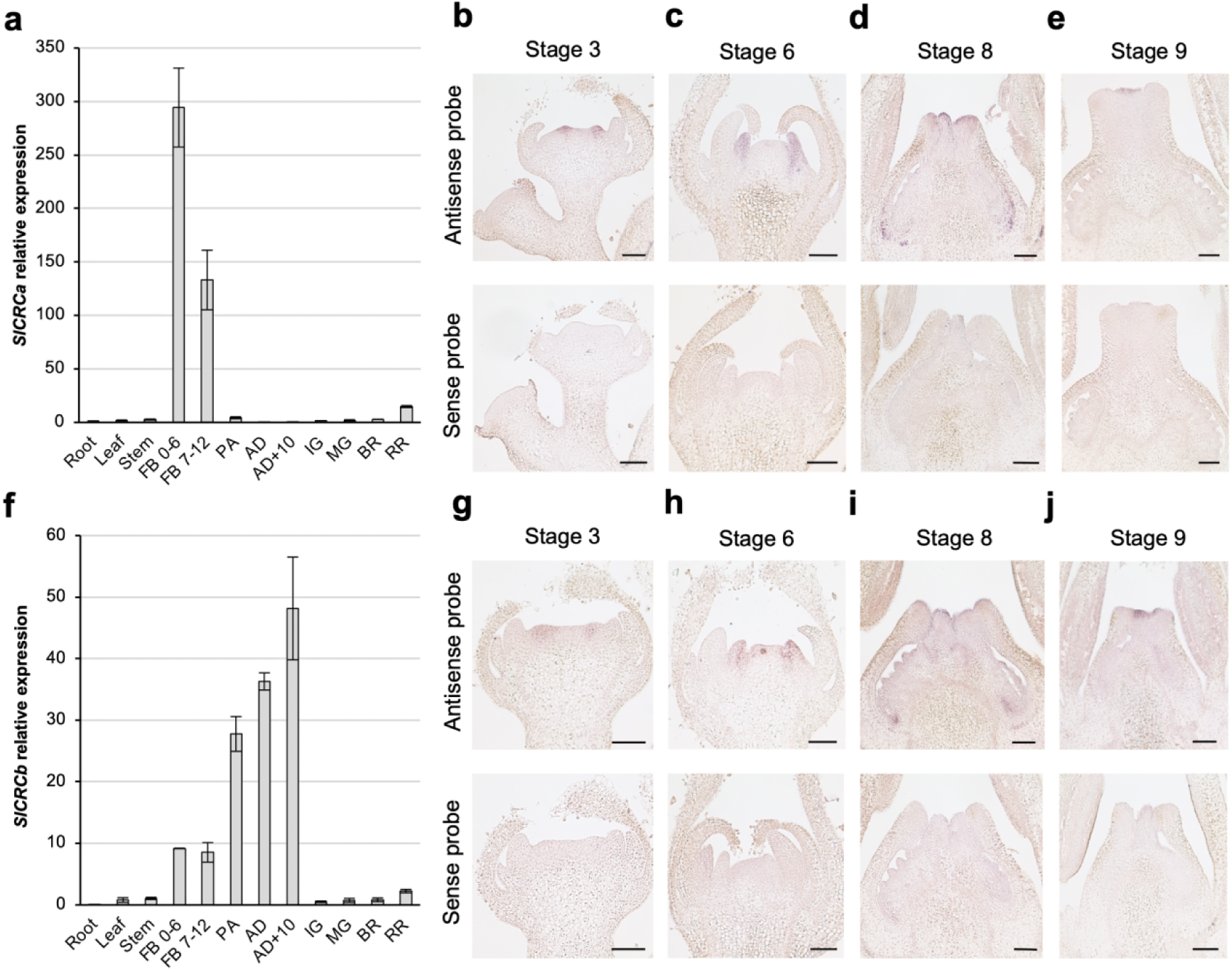
Dynamic expression of *SlCRCa* and *SlCRCb* genes. **a**, Relative expression of *SlCRCa* in different developmental tissues and stages of wild-type (WT) flowers. **b-e**, *In situ* mRNA hybridisation of *SlCRCa* using antisense or a sense probes in histological sections of WT flowers at different developmental stages: stage 3 (**b**), stage 6 (**c**), stage 8 (**d**) and stage 9 (**e**). **f**, Relative expression of *SlCRCb* in different developmental tissues and stages of WT flowers. **g-j**, *In situ* mRNA hybridisation of *SlCRCb* using antisense or a sense probes in histological sections of WT flowers at different developmental stages: stage 3 (**g**), stage 6 (**h**), stage 8 (**i**) and stage 9 (**j**). In **a**,**f** data are means ± standard deviations (SD) of three biological and two technical replicates. FB0-6, floral buds from stages 0 to 6; FB7-12, floral buds from stages 7 to 12; PA, flowers at pre-anthesis stage; AD, flowers at anthesis stages; AD+10, flowers 10 days after anthesis stage; IG; immature green fruit; MG, mature green fruit; BR, breaker fruit; RR, mature red fruit. In **b-e**,**g-j** scale bars represent 100 μm.

RNA sequencing (RNA-seq) was next performed on WT and *fig* floral buds at developmental stages 0-6 to gain insight into the functional role of *SlCRCa* during flower development. This analysis identified 2115 differentially expressed genes (DEGs) in the *fig* mutant as compared to WT (false discovery rate adjusted P-value < 0.05), in which 978 were up-regulated and 1137 were down-regulated (Supplementary Dataset 1), suggesting that decreasing *SlCRCa* expression considerably affected the transcriptome of *fig* floral buds. To investigate the functions of differentially expressed genes, we applied a Gene Ontology (GO) term enrichment analysis, which revealed 29 and 36 overrepresented GO terms for up- and down-regulated DEGs, respectively (Supplementary Dataset 2). For biological process, GO terms belonging to response to stimulus (GO:0050896) were enriched in both groups of DEGs. However, biological processes related to reproduction (GO:0000003), reproductive process (GO:0022414), developmental process involved in reproduction (GO:0003006), reproductive structure development (GO:0048608) and flower development (GO:0009908) were enriched among up-regulated DEGs. With regard to molecular function, GO terms involved in protein binding (GO:0005515) were enriched in both up- and down-regulated DEGs. In addition, transporter activity (GO:0005215) and transcription factor activity, sequence-specific DNA binding (GO:0003700) terms were enriched in up-regulated DEGs (Supplementary Dataset 2 and Supplementary Fig. 3).

Among the up-regulated DEGs annotated with the reproductive structure development (GO:0048608) and flower development (GO:0009908) terms, we found the tomato homologues of the Arabidopsis PIN-FORMED auxin efflux carrier (Solyc03g118740, Solyc04g007690 and Solyc05g008060) and the AUXIN RESPONSE FACTOR transcription factor (Solyc09g007810 and Solyc12g042070) families, as well as the tomato homologues of the floral homeotic genes *APETALA2* (*Solyc02g064960*), *APETALA3* (*Solyc04g081000*), *FRUITFULL* (*Solyc03g114830*) and *SEPALLATA4* (*Solyc03g114840*), the latter of which is also involved in the determination of FM (Ditta et al., 2004). Within this group of genes, we also found the homologue of the Arabidopsis *HECATE3* (*Solyc11g005780*), which encodes a bHLH transcription factor regulating the development of female reproductive tissues. Thus, the overexpression of *HECATE* genes causes the production of ectopic stigmatic tissue (Gremski et al., 2007). Remarkably, a second homologue of the Arabidopsis *CRC* gene (*SlCRCb, Solyc05g012050*) was moreover included in this group of up-regulated DEGs. In summary, RNA-seq analysis revealed that several genes involved in reproductive developmental process and gynoecium formation were affected by the lack of function of *SlCRCa* gene.

### Knockout mutations in *SlCRCb* mimic *fig* mutant phenotype

While the Arabidopsis genome carries one copy of the *CRC* gene, an ancestral duplication, which likely happened before the Solanaceae diversification, generated two paralogues, *CRCa/CRCb*, whose retention across Solanaceae genomes suggests functional relevance (Phukela et al., 2020). Sequence analysis of the Arabidopsis CRC (181 aa) and the tomato SlCRCa (173 aa) and SlCRCb (160 aa) putative amino acid sequences displayed that CRC has an identity of 64 and 68% to SlCRCa and SlCRCb, respectively; while SlCRCa shows 73% identity to the aminoacidic sequence of SlCRCb. Furthermore, the study of microsynteny conservation between genomic blocks harbouring *CRC* genes in Arabidopsis and tomato showed that the Arabidopsis genomic region containing *CRC* was more closely related to that from tomato which bears *SlCRCb* rather than *SlCRCa* (Supplementary Fig. 4). To gain further insight into the role of the *SlCRCb* gene, we examined the *SlCRCb* spatio-temporal expression by qRT-PCR and *in situ* hybridisation. As with *SlCRCa*, a specific expression in reproductive tissues was observed for *SlCRCb* (Fig. 3a,f), although *SlCRCb* transcripts were detected throughout all floral development stages, from floral buds at developmental stages 0-6 to flowers at 10 days post anthesis (Fig. 3f), whereas *SlCRCa* expression was mainly detected at first stages of the floral bud development (Fig. 3a). Even though *SlCRCa* and *SlCRCb* showed differential temporal expression patterns, *in situ* hybridisation studies in developing flowers revealed that *SlCRCb* shows an identical spatial expression profile to *SlCRCa*. Thus, *SlCRCb* transcript accumulation was first localised at floral stage 3 in carpel primordia (Fig. 3g), where it continued until floral stage 6 (Fig. 3h). As the development progressed to floral stage 8, *SlCRCb* expression was detected in the ovary walls and the distal regions of the developing gynoecium (Fig. 3i), where its expression remained restricted at floral stage 9 (Fig. 3j).

The incomplete penetrance and variable expressivity of mutations at the *SlCRCa* locus, as well as the overlapping *SlCRCa* and *SlCRCb* spatial expression patterns at the initiation of gynoecium development, led to hypothesised that a partial redundancy of tomato *CRC* paralogues may exist. To test this hypothesis, we generated knockout mutations by using CRISPR/Cas9 system with a single guide RNA targeting the second exon of the *SlCRCb* gene (Fig. 4a). We evaluated four independent T_0_ diploid lines (CR-*slcrcb*) homozygous or biallelic for edited knockout alleles (Fig. 4a). The CR-*slcrcb* lines produced flowers with ovaries composed of numerous carpels growing inside one another, which led to anomalous fruits similar to those observed in *fig* mutants, RNAi *SlCRCa* and CR-*slcrca* lines (Fig. 4b). CR-*slcrcb* lines also produced a wide diversity of flower and indeterminate fruit phenotypes, although the presence of fruits with weak or severe mutant phenotypes did not exceed 50%, being the percentages of fruits with severe mutant phenotype close to 15% (Fig. 4c). These results revealed that *SlCRCa* and *SlCRCb* could have partially redundant roles in FM determinacy.

**Fig. 4.**
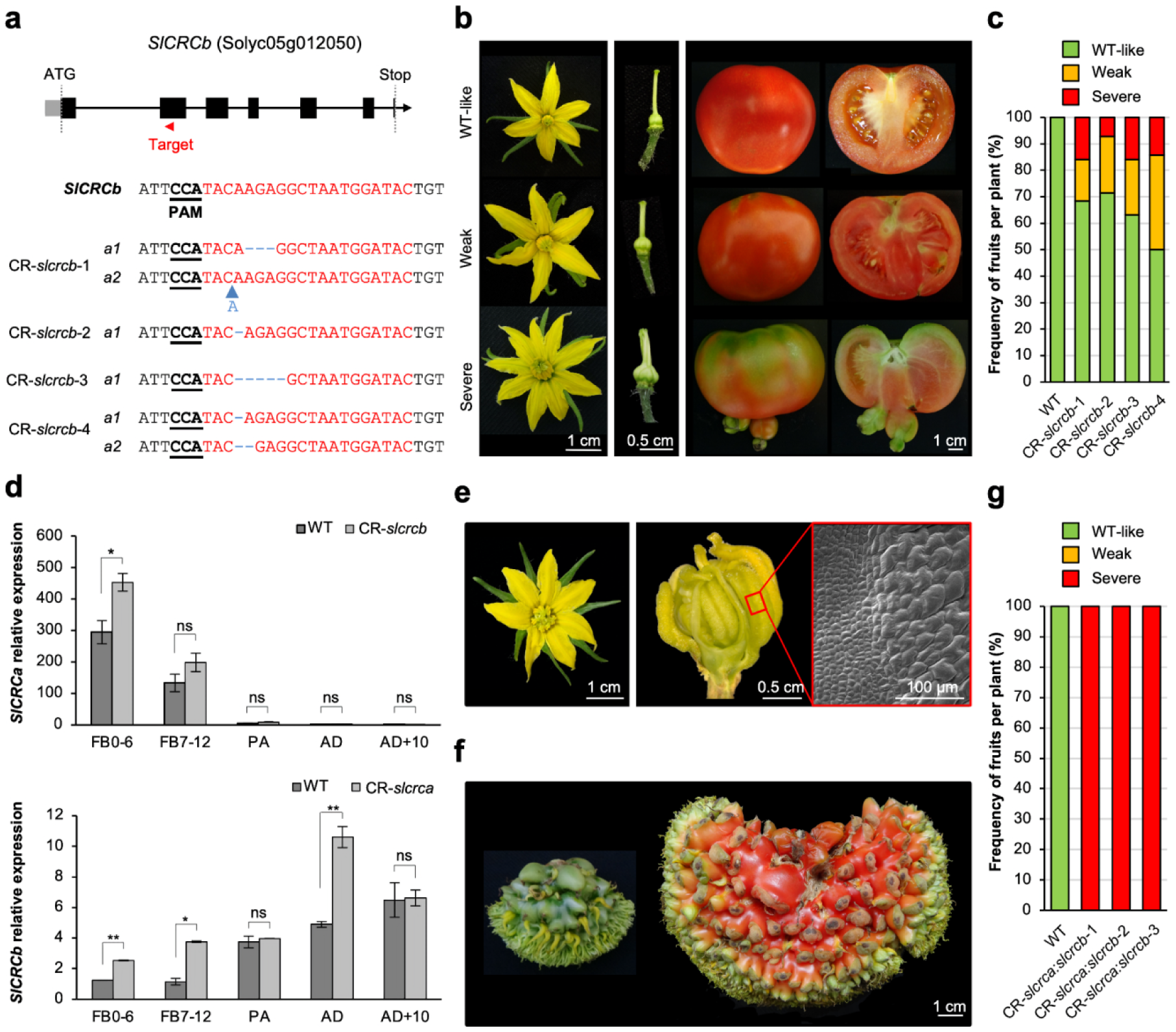
Characterisation of CRISPR/Cas9-*slcrcb* (CR-*slcrcb*) and double mutant CR-*slcrca:slcrcb* lines. **a**, CR-*slcrcb* alleles identified by cloning and sequencing PCR products from the *SlCRCb* targeted region from four T_0_ CRISPR plants. Black bold and underlined letters indicate protospacer-adjacent motif (PAM) sequences, blue dashed lines show InDel mutations, blue letter and arrow indicate an insertion sequence. **b**, Representative CR-*slcrcb* flowers, pistils and fruits exhibiting WT-like, weak and severe phenotypes. **c**, Percentage of different types of fruits harvested from wild-type (WT) and T_0_ CR-*slcrcb* plants. **d**, Relative expression of *SlCRCa* and *SlCRCb* in CR-*slcrcb* and CR-*slcrca* lines, respectively, at different floral developmental stages. FB0-6, floral buds from stages 0 to 6; FB7-12, floral buds from stages 7 to 12; PA, flowers at preanthesis stage; AD, flowers at anthesis stage; AD+10, flowers 10 days after anthesis stage. Data are means ± standard deviations (SD) of three biological and two technical replicates. A two-tailed, two-sample Student’s *t* test was performed, and significant differences are represented by asterisks: *, *P* < 0.01; **, *P* < 0.001; ns, no statistically significant differences. **e**,**f**, Representative flower, pistil and fruits developed by CR-*slcrca:slcrcb* double mutants. **e**, Detail of the fourth floral whorl organs and morphological features of their epidermal cells in a flower at anthesis stage. **f**, Immature green and mature red fruits. **g**, Percentage of different types of fruits harvested from WT and T_0_ CR-*slcrca:slcrcb* plants.

Taking into account that RNA-seq results showed that *SlCRCb* was up-regulated in *fig* floral buds, the question arose as to whether a compensatory mechanism between tomato *CRC* paralogues may be involved in gynoecium determination. For that purpose, the expression of *SlCRCa* and *SlCRCb* genes was quantified in reproductive tissues of *slcrcb* and *slcrca* CRISPR/Cas9 null mutants, respectively. *SlCRCb* was differentially up-regulated in CR-*slcrca* floral buds at developmental stages 0-6 (Fig. 4d). *SlCRCb* was also up-regulated in CR-*slcrca* from floral buds to anthesis day during floral development; whereas *SlCRCa* transcripts were up-regulated in CR-*slcrcb* floral buds at developmental stages 0-6 (Fig. 4d). Additionally, we generated CR-*slcrca:slcrcb* double mutant plants to further dissect the relationship between tomato *CRC* paralogues. Remarkably, the simultaneous loss of function of *SlCRCa* and *SlCRCb* resulted in homeotic alterations affecting carpel development as the shape of some of the cells of which they were composed attained a stamen-like nature (Fig. 4e and Supplementary Fig. 5). CR-*slcrca:slcrcb* flowers produced stamen-like carpels in a reiterating pattern exclusively affecting the fourth whorl, thus giving rise to fruits with a severe indeterminate phenotype (Fig. 4f,g), which in all cases was considerably more severe than in either CR-*slcrca or* CR-*slcrcb* single mutant plants. Hence, complete penetrance and invariable expressivity were found when both tomato paralogues lost their functions.

### Tomato CRC paralogues bind to the chromatin remodelling complex members repressing *SlWUS* expression

Floral determinacy requires the repression of the stem cell identity gene *SlWUS*^6^; therefore, we further investigated the function of tomato *CRC* paralogues in FM determination by examining the spatio-temporal expression of *SlWUS* in CR-*slcrc* lines by *in situ* hybridisation. In WT flowers, *SlWUS* expression in the organising centre was detected from early FM developmental stages (Chu et al., 2019), until stem cell activity was arrested in floral buds at stages 4-5, when the carpel primordia started to emerge (Fig. 5a). From stage 6 onwards, *SlWUS* signal was completely abolished in WT developing carpels. However, *SlWUS* transcripts were detected, between the growing but still unfused carpel primordia, in both CR-*slcrca* and CR-*slcrcb* floral buds at stage 6; and *SlWUS* expression was even observed at later stages in a small group of cells in the placenta, where the initiation of new carpel primordia probably occurs (Fig. 5a). As expected, an enlarged *SlWUS* expression domain was observed in CR-*slcrca:slcrcb* flowers leading to an increased FM size, which agrees with the severe indeterminacy found in double mutant carpels (Fig. 5a). By way of conclusion, the extended expression of *SlWUS* correlates with the floral indeterminacy phenotype of CR-*slcrc* lines and suggests that tomato *CRC* paralogues might participate in the regulation of *SlWUS* transcription limiting FM activity and promoting floral determinacy.

**Fig. 5.**
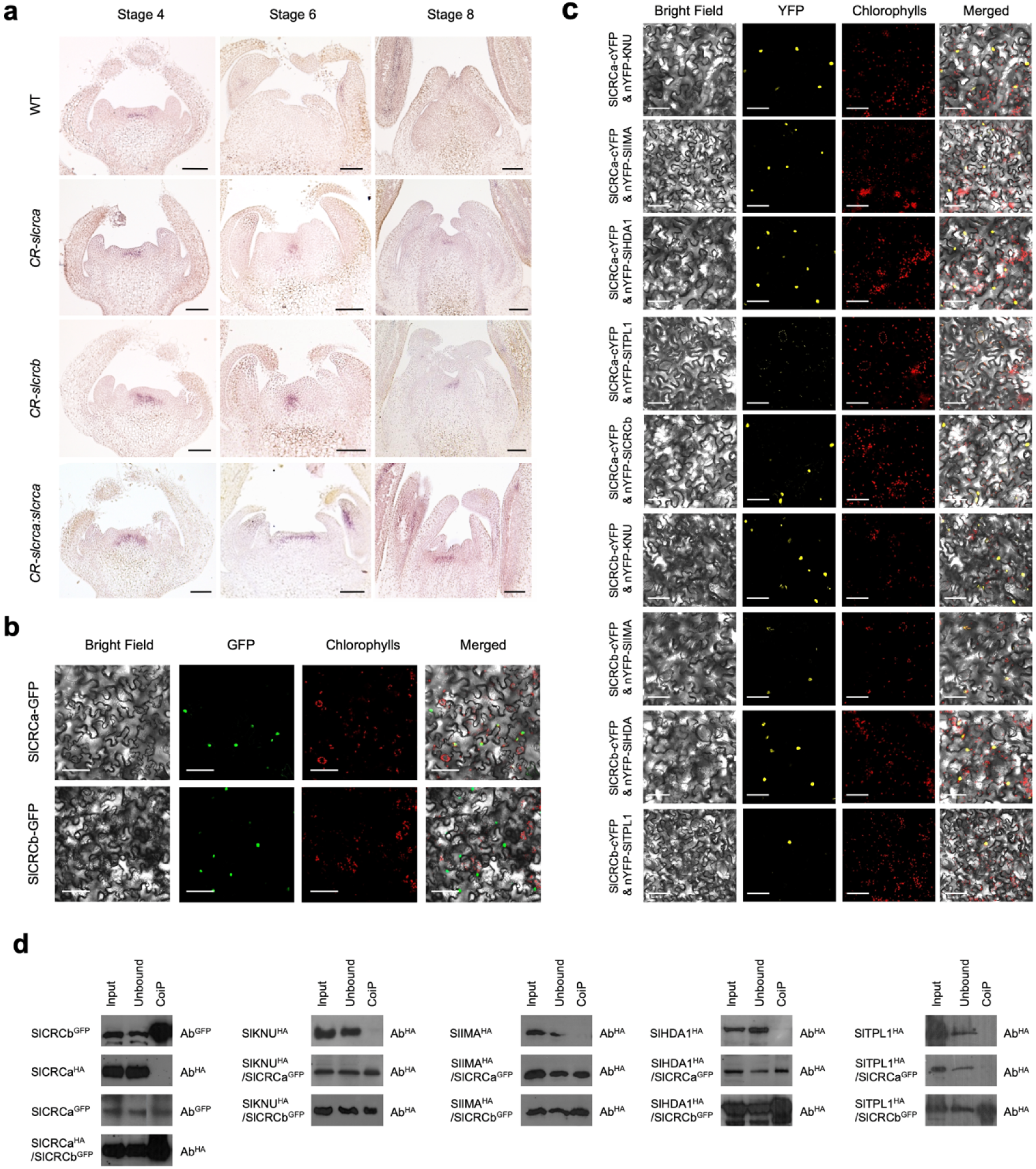
Tomato CRC paralogues interact with the chromatin remodelling complex members repressing *SlWUS* expression. **a**, *In situ* mRNA hybridisation of *SlWUS* in histological sections of wild-type (WT), CR-*slcrca*, CR-*slcrcb* and CR-*slcrca:slcrcb* flowers at developmental stages 4, 6 and 8. **b**, Subcellular localisation of SlCRCa and SlCRCb. The entire *SlCRCa* and *SlCRCb* coding sequences were N-terminal fused to green fluorescent protein (GFP) and transiently expressed in *Nicotiana benthamiana* leaves. **c**, Bimolecular fluorescence complementation confocal images showing *in vivo* interactions in *N. benthamiana* leaves between either the yellow fluorescent protein (YFP) C-terminal region fused to SlCRCa (SlCRCa-cYFP) or SlCRCb (SlCRCb-cYFP), and fusions of the YFP N-terminal region fused to SlKNU (nYFP-SlKNU), SlIMA (nYFP-SlIMA), SlHDA1 (nYFP-SlHDA1) or SlTPL1 (nYFP-SlTPL1). The SlCRCa-cYFP fusion was also examined with SlCRCb fused to the YFP N-terminal region (nYFP-SlCRCb). No YFP fluorescence signal was observed in leaves infiltrated with a single construct and the corresponding empty vector (Supplementary Fig. 6). **d**, Co-immunoprecipitation studies of *N. benthamiana* leaves expressing either GFP-tagged SlCRCa (SlCRCa^GFP^) or SlCRCb (SlCRCb^GFP^) and the different hemagglutinin (HA)-tagged members of the chromatin remodelling complex (SlKUN^HA^, SlIMA^HA^, SlHDA1^HA^ or SlTPL1^HA^). SlCRCb^GFP^ was also tested with HA-tagged SlCRCa (SlCRCa^HA^). The input total protein extracts were immunoprecipitated with anti-GFP beads, and the unbound and recovered fractions (CoIP) were incubated with anti-GFP (Ab^GFP^) and anti-HA (Ab^HA^) antibodies to detect precipitated and copurified proteins, respectively. In **a-c** scale bars represent 100 μm.

Recent studies have proven that the complete termination of floral stem cell activity in tomato is mediated by a chromatin remodelling complex, consisting of the SlIMA (Solyc02g087970), SlKNU, (Solyc02g160370) SlTPL (Solyc01g100050) and SlHDA1 (Solyc09g091440) proteins, which enables the complete repression of *SlWUS* (Bollier et al., 2018). The fact that *SlWUS* expression was mis-regulated in CR-*slcrca* and CR-*slcrcb* indeterminate flowers prompted us to question whether tomato *CRC* paralogues might be part of this regulatory pathway. Since SlKNU, SlIMA, SlTPL1 and SlHDA1 show nuclear localisation, we first evaluated SlCRCa and SlCRCb subcellular localisation by transient expression of N-terminal green fluorescent protein (GFP)-tagged versions of SlCRCa and SlCRCb. A confocal microscopy analysis revealed an exclusive nuclear localisation for both proteins (Fig. 5b). Next, we conducted bimolecular fluorescence complementation (BiFC) assays to evaluate whether tomato CRC paralogues might physically interact *in planta* with the chromatin remodelling complex members including SlKNU, SlIMA, SlTPL1 and SlHDA1. Thus, BiFC interactions were tested by transiently co-transforming with fusions of either the yellow fluorescent protein (YFP) C-terminal region fused to SlCRCa (SlCRCa-cYFP) or SlCRCb (SlCRCb-cYFP), and fusions of the YFP N-terminal region fused to either SlKNU (nYFP-SlKNU), SlIMA (nYFP-SlIMA), SlHDA1 (nYFP-SlHDA1) or SlTPL1 (nYFP-SlTPL1). In addition, the SlCRCa-cYFP fusion was co-transfected with SlCRCb fused to the YFP N-terminal region (nYFP-SlCRCb). As noted by the YFP fluorescent signal observed at the nucleus of epidermal cells, SlCRCa and SlCRCb were able to physically interact with each other and each of them with SlKNU, SlIMA, and SlHDA1, whereas SlCRCb was additionally found to bind to SlTPL1 (Fig. 5c). A lack of interaction between SlCRCa and SlTPL1 was also confirmed by BiFC experiments in the opposite orientation (Supplementary Fig. 6). To corroborate the interactions described above, they were furthermore tested by co-immunoprecipitation (CoIP) studies. Thus, co-transfection was performed with either GFP-tagged SlCRCa (SlCRCa^GFP^) or SlCRCb (SlCRCb^GFP^) and the different hemagglutinin (HA)-tagged members of the chromatin remodelling complex (SlKUN^HA^, SlIMA^HA^, SlHDA1^HA^ or SlTPL1^HA^). Likewise, the SlCRCb^GFP^ was co-transfected with HA-tagged SlCRCa (SlCRCa^HA^). Total protein extracts were inmunoprecipitated with anti-GFP beads. A Western blot analysis of the recovered fraction obtained using a specific antibody to the HA epitope revealed that SlCRCa^GFP^ copurified with SlKNU^HA^, SlIMA^HA^ and SlHDA1^HA^, but not with SlTPL1^HA^, as well as SlCRCb^GFP^ copurified with SlKNU^HA^, SlIMA^HA^, SlHDA1^HA^, SlTPL1^HA^ and SlCRCa^HA^ (Fig. 5d). Therefore, both BiFC and CoIP assays yielded concordant results, evidencing that SlCRCa and SlCRCb are able to physically bind to the chromatin remodelling complex members, which in turn repress *SlWUS* expression to promote FM determinacy.

### Tomato *CRC* paralogues partially rescue the loss-of-function of the Arabidopsis *CRC* gene

In Arabidopsis, mutations in *CRC* cause several phenotypic alterations affecting the gynoecium which fails to fuse at the apex, making it wider and shorter than the WT one (Alvarez and Smyth, 1999; Bowman and Smyth, 1999). As tomato *CRC* paralogues seem to have partially redundant functions, we next questioned whether they may be able to complement the phenotypic defects of Arabidopsis *crc* mutants. Thus, the Arabidopsis *crc-1* mutant, a strong hypomorphic allele in the ecotype Landsberg *erecta* (L*er*) genetic background, was genetically transformed with either *SlCRCa* or *SlCRCb* coding sequences under the control of the Arabidopsis *CRC* promoter (*pCRC*::*SlCRCa* and *pCRC*::*SlCRCb*). For these experiments, a 3860 bp fragment from upstream of the Arabidopsis *CRC* start codon was used as promoter sequence since its functionality has been previously demonstrated (Fourquin et al., 2007). In comparison to WT L*er* plants, *crc-1* mutants exhibited unfused carpels and the abolition of nectary development, as well as considerably shorter and wider siliques (Fig. 6). The *pCRC*::*SlCRCa* and *pCRC*::*SlCRCb* transgenes were able to fully restore carpel fusion (Fig. 6c), and slightly increased silique length (Fig. 6a,b). In agreement with the lack of nectaries in tomato flowers, *SlCRCa* and *SlCRCb* did not rescue nectary development in *crc-1* mutants (Fig. 6d). As a control, WT L*er* plants were also transformed with either *pCRC*::*SlCRCa* and *pCRC*::*SlCRCb*, which exhibited no significant differences with respect to untransformed L*er* plants (Supplementary Fig. 7). Overall, our results denote that tomato *CRC* paralogues are capable of partially restoring a WT phenotype when transformed into *crc-1* mutant plants.

**Fig. 6.**
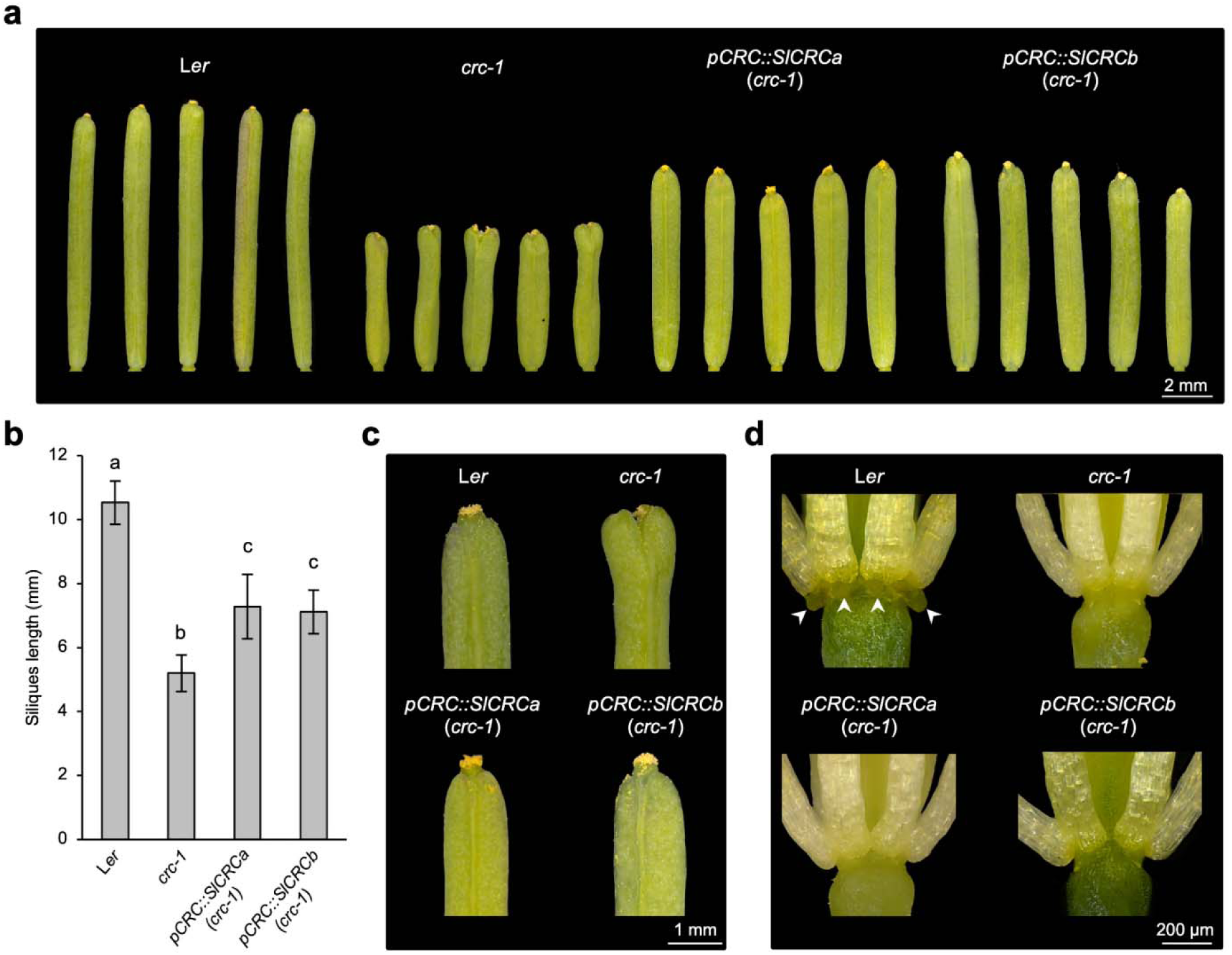
Complementation of the Arabidopsis *crc-1* mutation by transformation with tomato *CRC* paralogues. **a-d**, Fully elongated siliques (**a**), silique length (**b**), silique apices showing different degrees of carpel fusion (**c**), and development of nectaries (arrows) at the base of the third floral whorl in L*er* wild-type, which are absent in *crc-1* mutant and transgenic *pCRC*::*SlCRCa* and *pCRC*::*SlCRCb* plants (**d**). In **b** pairwise comparisons of means using the least significant difference test were performed. Values followed by the same letter (a, b, or c) are not statistically different (*P* < 0.05).

## Discussion

### Tomato *CRC* paralogues safeguard floral determinacy by acting in a partially redundant and compensatory manner

In the present study, we have addressed the functional characterization of tomato paralogous *CRC* genes and examined their potential roles in FM determinacy and carpel formation. Our results revealed that a 367 bp insertion in the fourth intron of the *SlCRCa* gene, a homologue of the Arabidopsis *CRC*, causes the carpel-inside-carpel phenotype observed in *fig* mutant plants. Loss-of-function analyses of *SlCRCa*, involving either knockdown (RNAi) or knockout (CRISPR/Cas9) approaches, allowed for the generation of an allelic series at this locus which resulted in analogous mutant phenotypes with incomplete penetrance and variable expressivity. Indeed, variable floral and fruit phenotypes were observed even in the same individual, ranging from indistinguishable from WT to severe developmental defects. Consequently, these results denoted that any impairment of *SlCRCa* function would give rise to a loss of FM determinacy characterized by the proliferation of additional carpels within the unfused primary carpels. Our results also suggested that there is not a direct correlation between *SlCRCa* mRNA levels and the severity of the mutant phenotypes, which agrees with the hypothesis that propose a non-linear gene dosage response for developmental regulators involved in complex transcriptional regulatory networks (Birchler et al., 2016). The lack of predictability between gene dosage and phenotypic alterations has been also reported for other tomato meristem genes. Thus, through creating a series of cis-regulatory alleles by genome editing, Rodríguez-Leal et al. (2017) demonstrated that variations in fruit size are not predicted by changes in gene expression levels of the CLAVATA-WUSCHEL signalling pathway.

Incomplete penetrance and variable expressivity of mutant phenotypes were also observed among knockout mutant lines for *SlCRCb*, whereas all flowers developed by double mutant plants for *SlCRCa* and *SlCRCb* loci showed a severe indeterminate phenotype. Based on these results, we conclude that tomato *CRC* paralogues operate as positive regulators of floral determinacy by acting in a partially redundant manner to safeguard normal determination of FM and carpel development. Indeed, the loss of carpel identity in CR-*slcrca:slcrcb* double mutant plants supports the role of tomato *CRC* paralogues in establishing carpel identity for proper completion of gynoecium and fruit developmental programs. Furthermore, *SlCRCa* was differentially up-regulated in tomato floral buds lacking *SlCRCb* function (CR-*slcrcb* plants), and vice versa, *SlCRCb* expression significantly increased in the absence of *SlCRCa* activity (CR-*slcrca* plants). Therefore, an active compensation mechanism of *SlCRCa* and *SlCRCb* functions may participate in the regulation of FM determinacy. In support of this hypothesis, transcriptional compensation has been recently described as a means to control meristematic activity in tomato, where the CLV3/embryo-surrounding region (CLE) ligand paralogues operate to buffer stem cell homeostasis (Rodriguez-Leal et al., 2019). Hence, the absence of *SlCRCa* or *SlCRCb* gene function would trigger an active compensation mechanism involving the upregulation of *SlCRCb* or *SlCRCa*, respectively, which help to buffer the severity of flower developmental alterations. However, environmental or other genetic factors could lead to a partially compensatory response influencing penetrance and expressivity of phenotypes associated with single mutations at either *SlCRCa* or *SlCRCb* loci. Taken together, our results indicate that tomato *CRC* paralogues operate as positive regulators of FM determinacy by acting in a partially redundant and compensatory manner to ensure normal floral development.

### Evolutionary conservation and divergence of *CRC* gene function in tomato

Based on functional studies, the role of putative *CRC* orthologues in FM determinacy and gynoecium development seems to have been conserved across angiosperms, including monocots, basal dicots and eudicots (Bowman and Smyth, 1999; Yamaguchi et al., 2004; Lee et al., 2005a, 2005b; Fourquin et al., 2005, 2007, 2014; Nakayama et al., 2010; Bartholmes et al., 2012; Sun et al., 2013; Morel et al., 2018). Nevertheless, specialised functions of *CRC*-like genes have been acquired after the evolutionary divergence of their respective plant lineages. Thus, in monocot species such as rice, the *CRC* orthologue *DROOPING LEAF* (*DL*) is also involved in carpel organ identity and plays an essential role in leaf midrib formation, functions that are shared by *CRC* orthologues from other monocot species (Yamaguchi et al., 2004; Ishikawa et al., 2009; Wang et al., 2009; Nakayama et al., 2010; Strable et al., 2017). Likewise, novel functions in stigmatic cavity formation and ovule initiation have been proposed for the paralogous *CRC* genes *PeDL1* and *PeDL2* of the orchid *Phalaenopsis equestri* (Chen et al., 2021). Within the Solanaceae family, the functional role of *CRC* paralogues has also been addressed in petunia, where *PhCRC1* and *PhCRC2* genes, in addition to regulating floral determinacy and carpel development, are required for nectary development acting in a redundant manner to trigger its formation (Morel et al., 2018). *PhCRC1* and *PhCRC2* showed quasi-identical expression profiles, which displayed an accumulation of their transcripts in carpel primordia, ovary walls, and style and stigma (Morel et al., 2018), similar to the pattern observed for the tomato *CRC* paralogues at the beginning of flower development. However, despite the fact that tomato *CRC* paralogues play partially redundant roles to ensure normal floral development, they were differentially expressed during flower ontogenesis. Thus, *SlCRCa* was mainly expressed at the first stage of the flower bud formation, whereas *SlCRCb* transcripts were detected from young floral buds to flowers at 10 days post anthesis, suggesting that their regulatory elements may have diverged. In summary, although fine-tuning gene regulation events may have favoured gene paralogue speciation, flower and fruit phenotypes of tomato plants lacking *SlCRCa* and *SlCRCb*, together with gene expression analyses provide sufficient evidence about the conserved function of these genes in regulating carpel development and floral determinacy. Indeed, the heterologous expression of either *SlCRCa* or *SlCRCb* genes in Arabidopsis *crc-1* background was capable of partially restoring a WT phenotype, supporting the evolutionary ancestral role of *CRC*-like genes in promoting floral stem cell termination and carpel formation. However, tomato *CRC* paralogues were not able to rescue the proper formation of nectaries. Previous studies have reported that *CRC* gene functions in nectary development appears to be conserved in several core eudicot species, whereas *CRC*-like genes are not required for nectary formation in basal angiosperms, which would support the hypothesis that *CRC* function in nectary specification may be the consequence of *CRC* neofunctionalization within diverse lineages of core eudicots (Lee et al., 2005b; Fourquin et al., 2005, 2014; Yamada et al., 2011).

In Arabidopsis and rice, *CRC* orthologue genes have also been found to act antagonistically with class B genes in promoting carpel development (Alvarez and Smyth, 1999; Bowman and Smyth, 1999; Yamaguchi et al., 2004). Here, we first describe the *CRC* functional role in tomato, and we found that double mutant plants for *SlCRCa* and *SlCRCb* loci develop flowers with stamen-like carpels growing in a reiterating pattern inside the fourth whorl, which strongly hints that tomato CRC paralogues seem to have common functions with the Arabidopsis and rice counterparts as negative regulators of class B genes. Accordingly, a recent research in the Solanaceae family member *Physalis floridana,* has revealed regulatory and genetic interactions between B-class MADS-box and *CRC* genes in a context-dependent manner during flower development (Gong et al., 2021).

### Tomato CRC paralogues are part of the chromatin remodelling complex that repress *SlWUS* in floral meristems

A complex regulatory network involving signalling cascades, transcriptional regulation, epigenetic mechanisms and hormonal control for FM determinacy has been described in the model species Arabidopsis (Shang et al., 2019). Thus, the function of *CRC* as regulator of floral determinacy has hitherto been limited to modulate auxin homeostasis by both activating *YUC4* and repressing *TRN2* gene expression (Yamaguchi et al., 2017, 2018). Our findings reveal a new role for tomato *CRC* paralogues in balancing floral stem cell proliferation and differentiation, as they are able to physically bind to the members of the chromatin remodelling complex that drives the epigenetic regulation of *SlWUS* expression. In this epigenetic silencing mechanism, SlIMA acts as an adaptor protein engaging SlKNU in a complex that involves SlTPL and SlHDA1 leading to *SlWUS* repression (Bollier et al., 2018). Based on the *SlWUS* expression profiles in flowers of single and double mutants for *SlCRCa* and *SlCRCb*, as well as protein interaction data hereby reported, it seems reasonable to propose a model by which *SlCRCa* and *SlCRCb* would act in specifying floral determinacy by binding to the chromatin remodelling complex that ensures the proper spatio-temporal repression of *SlWUS* during flower development (Fig. 7). A future challenge will be to assess the role of tomato *CRC* paralogues in regulating auxin homeostasis, as well as to determine whether CRC interactions are conserved in angiosperm species. Further studies on the degree of conservation or divergence in the molecular mechanism triggering floral determinacy will provide valuable information for crop yield improvement, as the number of carpels in a flower, and consequently the final number of locules forming the mature fruit, plays a key role in regulating fruit size. In conclusion, this research contributes to a better understanding of the genetic and molecular mechanisms regulating FM determinacy, and hence influencing external fruit quality, an important trait in tomato breeding. As it has been proven for the CLAVATA-WUSCHEL signalling network (Rodríguez-Leal et al., 2017; Liu et al., 2021), the ever-increasing understanding of meristem regulatory pathways, and their relevance in regulating crop yield, allow for new knowledge-driven breeding strategies, which will in turn significantly contribute to the sustainability of agriculture in the next decades.

**Fig. 7.**
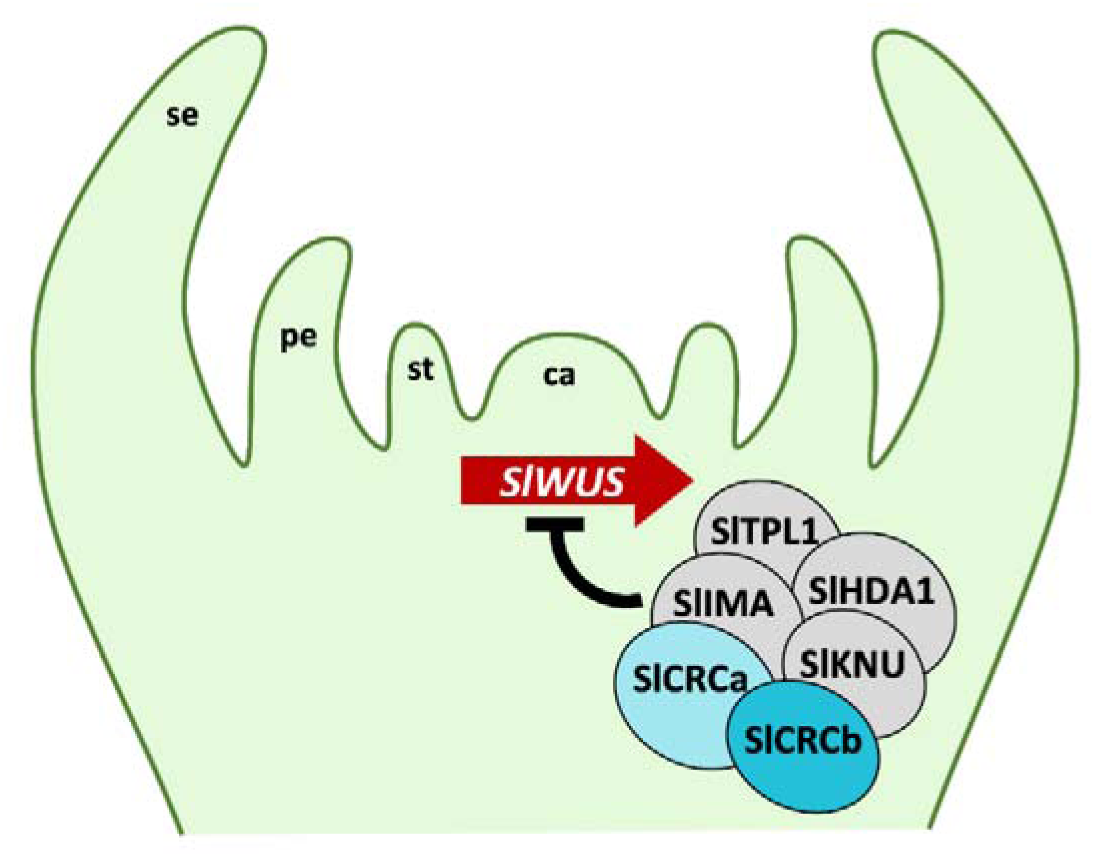
Proposed model for the function of tomato CRC paralogues in *SlWUS* repression and floral meristem determinacy. Tomato CRC paralogues (SlCRCa and SlCRCb), SlIMA and SlKNU interact with SlHDA1 and SlTPL1 to form a chromatin remodelling complex that represses *SlWUS* expression to terminate floral stem cell activity once carpel primordia are initiated. se, sepal; pe, petal; st, stamen; ca, carpel.

## Methods

### Plant material and growth conditions

The *fig* mutant was identified from a T-DNA insertional mutant collection generated in the genetic background cultivar P73 (Pérez-Martín et al., 2017). However, molecular analysis showed that the *fig* mutation was not associated with a T-DNA insertion (Supplementary Fig. 2). For mapping-by-sequencing, an F_2_ mapping population was generated by crossing a *fig* mutant to wild tomato *Solanum pimpinellifolium* accession LA1589 and self-fertilizing the F_1_ plants. Tomato plants were grown under greenhouse conditions with natural sunlight (13 to 14 h photoperiod). Temperatures ranged between 26-34°C and 20-25°C for daytime and night-time periods, respectively. Standard management practices were used including regular addition of fertilisers.

All *Arabidopsis thaliana* plants used for this study, including mutant and transgenic plants, were in the ecotype Landsberg *erecta* (L*er*) genetic background (kindly provided by Prof. M.R. Ponce, Miguel Hernández University, Elche, Spain). Seeds of the *crabs claw-1* (*crc-*1; CS3814; N3814) mutant (Alvarez and Smyth, 1999) were initially obtained from the Nottingham Arabidopsis Stock Center (NASC; Nottingham, UK) and propagated at our laboratory for further analysis. Seeds were surface sterilised for 8 min in 40% (v/v) sodium hypochlorite and 1% (v/v) Triton X-100 with periodic agitation, rinsed at least three times with sterile distilled water and sown in Petri dishes containing MS (Murashige and Skoog, 1962) growth medium for germination. When required, kanamycin was added to the MS growth medium at a final concentration of 50 μg/ml. After stratification at 4°C for 24 h in the dark, the plates were incubated in a growth chamber at 22°C with a 16 h light / 8 h darkness cycle. For non-sterile conditions, plants were grown in pots at 25°C and 60–70% relative humidity under constant illumination.

### Phenotypic characterisation of tomato flowers and fruits

The number of floral organs (sepals, petals, stamens, and carpels) was evaluated in at least 60 flowers at anthesis stage (AD) showing WT-like, weak and severe phenotypes. A minimum of 60 mature fruits were collected and used to calculate the average fruit weight (g), width (mm), length (mm) and number of locules per phenotype. All values were expressed as the mean ± standard deviation. Significant differences were calculated by pairwise comparisons of means using the least significant difference test.

For optical microscopy analysis, floral buds were harvested at different developmental stages. Samples were fixed in FAE (10% formaldehyde, 5% acetic acid, and 50% absolute ethanol), dehydrated in ethanol series, embedded in paraffin and cut using a Leica RM2035 microtome. Eight µm transversal sections were stained in a 1% Toluidine Blue solution and observed under a Leica DM6 B microscope. Scanning electron microscopy analyses were performed as described in Lozano et al. (1998). Floral bud samples were hand-dissected, fixed in FAEG (10% formaldehyde, 5% acetic acid, 50% absolute ethanol and 0.72% glutaraldehyde) and dehydrated in ethanol series. Subsequently, tissues were dried using a critical point dryer (Bal-Tec CPD 030) and gold coated in a Sputter Coater (Bal Tec SCD005). Finally, samples were visualised using a Hitachi S-3500N scanning electron microscope at 10 kV.

### Whole genome sequencing and allele frequency analysis

Mapping-by-sequencing was performed as described previously Yuste-Lisbona et al. (2021). Briefly, two pools were formed using equal amounts of DNA from 25 mutant and 50 randomly selected wild-type plants. DNA quality control, Illumina library construction and sequencing with 100 bp paired-end reads on the Illumina HiSeq2000 platform (Illumina, Inc., San Diego, CA) were carried out at the Genome Center of the Max Planck Institute for Plant Breeding Research (Cologne). The resulting short reads have been deposited in the Sequence Read Archive (SRA) database at the National Center for Biotechnology Information (NCBI) under BioProject accession number PRJNA685617. Paired-end reads were aligned to the tomato genome reference sequence version 4.0 (ITAG4.0) using Bowtie2 version 2-2.0.0-b5 with default parameters (Langmead and Salzberg, 2012). Duplicated were removed with MarkDuplicates from the Picard tools (http://broadinstitute.github.io/picard/). Variant calling analysis was carried out using the HaplotypeCaller tool from GATK (DePristo et al., 2011). Next, BCFtools from the SAMtools package (Li et al., 2009) was used to filter biallelic SNPs with a minimum depth of 10 reads in each sample. After filtering, the allele frequency ratio (i.e., non-reference allele counts / total allele counts) for biallelic variants was calculated. Finally, to identify the chromosomal region where the *fig* mutation is located, the average allele frequencies were plotted along each chromosome using a custom script in the R environment for statistical computing (R Development Core Team, 2020) which uses a sliding window and step size of 1000 and 100 variants, respectively. Once the candidate genomic region to host the *fig* mutation was determined, the identified variants encompassing this genomic region were filtered based on the following criteria: i) variants that were heterozygous (0/1) and homozygous (1/1) for the alternative allele in wild-type and mutant pools, respectively; and ii) unique variants that were not previously reported in the sequenced tomato genomes (Lin et al., 2014; The 100 Tomato Genome Sequencing Consortium, 2014).

### PCR genotyping of *SlCRCa* locus

DNAzol® Reagent kit (Invitrogen Life Technologies, San Diego, CA, USA Genomic DNA) was used to extract genomic DNA from young leaves of a segregating *fig* line following the manufacturer’s instructions. DNA quantification was carried out using a NanoDrop 2000 spectrophotometer (Thermo Scientific, Wilmington, DE, USA). *SlCRCa* locus was genotyped using the SlCRCa-Fg and SlCRCa-Rg primers which amplify a 332 bp fragment in wild-type homozygous plants, a 699 bp fragment in *fig* mutant plants and both of them in heterozygous ones. The sequences of primers used are shown in Supplementary Table 2.

### RNA isolation and Quantitative Real-Time PCR

Total RNA was extracted using TRIZOL (Invitrogen Life Technologies, San Diego, CA) according to the manufacturer’s indications. RNA quantity was estimated using a NanoDrop 2000 spectrophotometer (Thermo Scientific, Wilmington, DE, USA) and quality was checked by gel electrophoresis. Contaminating DNA was removed using the DNA-freeTM kit (Ambion, Austin, TX, USA) and 1 µg of RNA was used for cDNA synthesis with a ML-MLV reverse transcriptase (Invitrogen) with a mixture of random hexamer and oligo-(dT)_18_ primers. The qRT-PCR analysis was carried out using three biological and two technical replicates. Specific primer pairs were used in each qRT-PCR reaction with the SYBR Green PCR Master Mix kit (Applied Biosystems, Foster City, CA, USA) on the 7300 Real-Time PCR System (Applied Biosystems, Foster City, CA, USA). The housekeeping gene *Ubiquitin3* (*Solyc01g056940*) was used as a control in all gene expression analyses. Quantification of gene expression was performed using the ΔΔCt calculation method (Winer et al., 1999). Relative expression values were represented as the mean ± standard deviation. The sequences of primers used in this study are listed in Supplementary Table 3.

### Generation of tomato transgenic lines

To generate the RNA interference (RNAi) *SlCRCa* construct, a 118 bp fragment of the first exon of the *Solyc01g0104120* cDNA was amplified using the SlCRCa-Fi and SlCRCa-Ri primers and cloned in sense and antisense orientation into the pKannibal vector (Wesley et al., 2001). The modified pKannibal vector was digested with *Not*I and the resulting restriction fragment was cloned into the pART27 binary vector (Gleave, 1992). CRISPR-Cas9 lines of *SlCRCa* and *SlCRCb* genes were obtained following the protocol described by Vazquez-Vilar et al. (2016). Thus, Breaking-Cas web software (Oliveros et al., 2016) was used to design the sgRNA target sequences within the coding region of each *SlCRC* gene, *SlCRCa* (GTATCCAACAACTTCTTGCA) and *SlCRCb* (GTATCCATTAGCCTCTTGTA). Each first-generation (T_0_) transgenic plant was genotyped with primers that cover the target recognition region of each sgRNA. PCR products were cloned into the pGEM-T vector and at least 10 clones per PCR product were sequenced. Finally, only biallelic or chimeric T_0_ plants for mutant knockout alleles were phenotyped. The sequences of primers used in the generation of RNAi and CRISPR/Cas9 constructs are shown in Supplementary Table 2. Genetic transformation experiments were carried out as described in Ellul et al. (2003). using *Agrobacterium tumefaciens* LBA4404 strain. The ploidy levels of T_0_ transgenic plants were assessed by following as previously described by Atarés et al. (2011). The CR-*slcrca:slcrcb* double mutant plants were generated using standard crossing between single *slcrcb* and *slcrca* T_0_ CRISPR null mutants, and were confirmed by genotyping from F_1_ progeny plants.

### *In situ* hybridisation analysis

Tissue preparation, sectioning and transcript detection for *in situ* hybridisation experiments were carried out as described in Lozano et al. (1998). Briefly, *SlCRCa* (*Solyc01g0104120*), *SlCRCb* (*Solyc05g012050*) and *SlWUS* (*Solyc11g071380*) probes were synthesised using cDNA as template by using the primers SlCRCa-Fz/SlCRCa-Rz, SlCRCb-Fz/SlCRCb-Rz and SlWUS-Fz/SlWUS-Rz (sequences are reported in Supplementary Table 3), respectively, and the resulting products were cloned into the pGEM-T Easy vector (Promega, Madison, WI, USA). After plasmid linearisation, the DIG RNA labelling mix (Roche Applied Science, Indianapolis, IN, USA) and, depending on the orientation of the insert, T7 and SP6 polymerases were used for *in vitro* transcription of antisense probes. As negative control, sense RNA probes were synthesised and hybridised to sections of tomato floral buds.

### RNA sequencing

For expression analyses, three biological replicates per genotype were sequenced, each with at least 30 floral buds at developmental stages 0-6. Total RNA was isolated using TRIZOL (Invitrogen Life Technologies, San Diego, CA) following the manufacturer’s instructions. Contaminating DNA was removed using the DNA-free^TM^ kit (Ambion, Austin, TX, USA). Libraries were prepared according to the Illumina TruSeq RNA protocol and sequenced with paired-end 150 bp on the Illumina HiSeq2000 platform (Illumina, Inc., San Diego, CA, USA). The resulting short reads were deposited in the SRA database at the NCBI under BioProject accession number PRJNA686085. The sequence reads were aligned to the tomato genome reference sequence version 4.0 (ITAG4.0) using Tophat v2.0.6 (Kin et al., 2013) with the following parameters: --max-insertion-length 12 --max-deletion-length 12 -g 1 --read-gap-length 12 --read-edit-dist 20 --read-mismatches 12 --no-coverage-search --read-realign-edit-dist 0 --segment-mismatches 3 --splice-mismatches 1. The raw number of reads per transcript was counted using the Bioconductor packages GenomicFeatures and GenomicAlignments (Lawrence et al., 2013). Differentially expressed genes were determined using the Wald test in the DEseq2 package (Love et al., 2014) with the Benjamini-Hochberg multiple testing correction (Benjamini and Hochberg, 1995) for false discovery rate (FDR). Genes with an FDR adjusted P-value of less than 0.05 were defined as significantly up- or down-regulated. Gene Ontology (GO) term enrichment analysis of the significantly differentially expressed genes was performed using agriGO v2.0 software (Tian et al., 2017) and determined by FDR < 0.05 with the Fisher statistical test and the Bonferroni multi-test adjustment.

### Molecular complementation of the Arabidopsis *crc-1* mutant

The Arabidopsis *crc-1* mutant was complemented with two different constructs, each carrying a 3860 bp fragment from upstream of the Arabidopsis *CRC* start codon representing its promoter region (*pCRC*), fused to the coding sequences of either the tomato *SlCRCa* (*pCRC::SlCRCa*) or *SlCRCb* (*pCRC::SlCRCb*) genes. The primers used for PCR amplification of *pCRC* and genes coding regions are shown in Supplementary Table 4. PCR products were purified and cloned into the pGEMT vector (Promega, Madison, WI, USA). *pCRC* and each gene coding fragments were linked by a double digestion with *Xho*I and a specific restriction enzyme which cuts within the pGEM-T polylinker (*Aat*II or *Sac*I depending on the *pCRC* and coding sequences orientation in the pGEM-T) and subsequent ligation conducted by T4 DNA ligase. The complete sequences of the *pCRC* fused to the corresponding gene coding sequence were obtained by amplifying with pCRC-Fac/SlCRCa-Rac or pCRC-Fac/SlCRCb-Rac primers (Supplementary Table 4), then cloned into the pENTR/D-TOPO vector (Invitrogen Life Technologies, San Diego, CA, USA), and finally subcloned into the Gateway binary vector pGWB401 (Nakagawa et al., 2007). All final plasmids were verified by Sanger sequencing and then transformed into *A. tumefaciens* C58C1 strain. The control WT L*er* and the mutant *crc-1* plants were transformed by the floral dip method described by Clough and Bent (1998). T_1_ transgenic plants were selected on plates supplemented with 50 μg/mL kanamycin as previously described in Harrison et al. (2006). At least 30 flowers and siliques from L*er*, *crc-1* and T_1_ transgenic plants, resulting from the transformation of L*er* and *crc-1* plants with either *pCRC*::*SlCRCa* or *pCRC*::*SlCRCb* constructs, were evaluated on a Leica DMS1000 digital microscope.

### Sequence alignment and microsynteny analysis

The amino acid sequences of SlCRCa (XP_004228849), SlCRCb (XP_004239032) and CRC (NP_177078) were downloaded from the GenBank database (http://www.ncbi.nlm.nih.gov) and pairwise aligned using BLASTp (https://blast.ncbi.nlm.nih.gov/Blast.cgi). The study of microsynteny conservation between the genomic regions harbouring the Arabidopsis *CRC* gene and either the tomato *SlCRCa* or *SlCRCb* genes was carried out with the GEvo tool available in the CoGe server (https://genomevolution.org/coge/GEvo.pl). Thus, pairwise genomic alignments were performed between the genomic region comprising 10,000 nt upstream and downstream of the Arabidopsis *CRC* gene (chromosome 1 positions: 25,997,465-26,019,059; TAIR v10.02), and either 30,000 nt of the genomic region flanking the tomato *SlCRCa* (chromosome 1 positions: 5,022,049-5,084,644; ITAG4.0) or *SlCRCb* (chromosome 5 positions: 5,323,844-5,385,856; ITAG4.0) genes.

### Subcellular localisation and bimolecular fluorescence complementation assays

SlCRCa and SlCRCb subcellular localisations were assessed by fusing each protein to the green fluorescent protein (GFP). Thus, full-length open reading frame sequences coding for SlCRCa and SlCRCb proteins were cloned into the pENTR/D-TOPO vector (Invitrogen Life Technologies, San Diego, CA, USA) and recombined into the Gateway binary vector pGWB6 (Nakagawa et al., 2007) to integrate GFP at the N–terminus of the proteins of interest. To test for bimolecular fluorescence complementation (BiFC)-based protein–protein interaction, full-length open reading frame sequences of SlCRCa, SlCRCb, SlKNU, SlIMA, SlHDA1 and SlTPL1 proteins were cloned into the pENTR/D-TOPO vector and subloned into the Gateway binary vectors containing the N- or C-terminal fragments of the yellow fluorescent protein (pYFN43 and pYFC43 vectors, respectively). Constructs were transformed in *A. tumefaciens* GV3101 strain and infiltrated into *Nicothiana benthamiana* leaves from 2-3-week-old plants. Plants were kept in long-day (16 h light/8 h dark) conditions at 22°C. Samples were observed 3 days post infiltration using a Nikon ECLIPSE Ti confocal microscope equipped with Nikon C2 Si lasers. The sequences of primers used for subcellular localisation and BiFC assays are shown in Supplementary Table 5.

### Co-immunoprecipitation assays

For co-immunoprecipitation assays, *N. benthamiana* leaves were transiently co-transfected with *A. tumefaciens* GV3101 strain cultures expressing either GFP-tagged SlCRCa (SlCRCa^GFP^) or SlCRCb (SlCRCb^GFP^) and the different hemagglutinin (HA)-tagged members of the chromatin remodelling complex (SlKUN^HA^, SlIMA^HA^, SlHDA1^HA^ or SlTPL1^HA^). Likewise, the SlCRCb^GFP^ was co-transfected with HA-tagged SlCRCa (SlCRCa^HA^). For this purpose, pENTR/D-TOPO vectors containing full-open reading frame sequences of the recombinant fusion proteins of interest were recombined into the Gateway binary vectors pGWB6 and pGWB15 (Nakagawa et al., 2007), containing GFP or HA tags, respectively. These vectors were co-infiltrated in *N. benthamiana* leaves from 2-3-week-old plants. Plants were kept under long-day (16 h light/8 h dark) conditions at 22°C. Subsequent protein extraction was performed from leaves harvested 2 days after infiltration. Plant material was ground in liquid nitrogen and homogenised in protein extraction buffer (25 mM Tris-HCl, pH 7.4, 75 mM NaCl, 0.5% Nonidet P-40, 0.05% sodium deoxycholate, 10 mM β-mercaptoethanol, 1 mM PMSF, and cOmplete-EDTA free protease inhibitor cocktail (Roche Applied Science, Indianapolis, IN, USA). Protein extracts were centrifuged twice at 14,000 g for 10 min at 4°C. After cell lysis, GFP-tagged proteins were magnetically labelled and subsequent isolated using µMACS Isolation Kit (Miltenyi Biotec S.L., Bergisch Gladbach, Germany) following manufacturer’s instructions. The resulting samples were analysed by SDS-PAGE and immunoblotted using anti-GFP-HPR (Miltenyi Biotec, Bergisch Gladbach, Germany) and anti-HA-peroxidase (Roche Applied Science, Basel, Switzerland) antibodies.

### Accession numbers

Tomato sequence data from this article can be found at the SOL Genomics Network (SGN; http://sgn.cornell.edu) under accession numbers Solyc01g010240 (*SlCRCa*), Solyc05g012050 (*SlCRCb*), Solyc11g071380 (*SlWUS*), Solyc02g087970 (*SlIMA*), Solyc02g160370 (*SlKNU*), Solyc01g100050 (*SlTPL*), and Solyc09g091440 (*SlHDA1*). Sequence data from Arabidopsis can be found at the Arabidopsis Information Resource (TAIR; https://www.arabidopsis.org/) under the accession number At1g69180 (*CRC*). The DNA-seq and RNA-seq data from this article can be found at the Sequence Read Archive (SRA; https://www.ncbi.nlm.nih.gov/sra/) under BioProject accession numbers PRJNA685617 and PRJNA686085, respectively.

## Acknowledgments

The authors thank Cristina Ferrándiz for critical review of the manuscript. This work was supported by the PID2019-110833RB-C31 and PID2019-110833RB-C32 grants from the Spanish Ministry of Science and Innovation (MICI/AEI/FEDER, UE) and the BRESOV (Breeding for Resilient, Efficient and Sustainable Organic Vegetable production) project funding by the Research and Innovation Programme of the European Union Horizon 2020 (grant agreement No. 774244). The authors also thank research facilities provided by the Campus de Excelencia Internacional Agroalimentario (CeiA3).

## Author contributions

L.C. performed most experimental procedures, prepared figures and co-wrote the manuscript. E.G., B.P., B.G.-S., A.O.-A. and R.M.-P. participated in the experimental research. T.A., J.C., V.M., F.J.Y.-L., and R.L. contributed materials and genetic resources, and analysed data. F.J.Y.-L., and R.L. conceived and supervised research and co-wrote the paper.

